# Optimal linear estimation models predict 1400-2800 years of co-existence between Neandertals and *Homo sapiens* in western Europe

**DOI:** 10.1101/2022.06.20.496862

**Authors:** Igor Djakovic, Alastair Key, Marie Soressi

## Abstract

Recent fossil discoveries suggest that Neandertals and *Homo sapiens* may have co-existed in Europe for as long as five to six thousand years. Yet, evidence for their contemporaneity at any regional scale remains elusive. In France and northern Spain, a region which features some of the latest directly-dated Neandertals in Europe, Protoaurignacian assemblages attributed to *Homo sapiens* appear to ‘replace’ Neandertal-associated Châtelperronian assemblages. Using the earliest and latest *known* occurrences as starting points, Bayesian modelling has provided some indication that these occupations may in fact have been partly contemporaneous. The reality, however, is that we are unlikely to ever identify the ‘first’ or ‘last’ appearance of a species or cultural tradition in the archaeological and fossil record. Here, we use optimal linear estimation modelling to estimate the first appearance date of *Homo sapiens* and the extinction date of Neandertals in France and northern Spain by statistically inferring these ‘missing’ portions of the Protoaurignacian and Châtelperronian archaeological records. Additionally, we estimate the extinction date of Neandertals in this region using a set of directly-dated Neandertal fossil remains. The results suggest that the onset of the *Homo sapiens* occupation of this region likely preceded the extinction of Neandertals and the Châtelperronian by up to 1400-2900 years – raising the possibility of an extended co-existence of these groups during the initial Upper Palaeolithic of this region. Whether or not this co-existence featured some form of direct interaction, however, remains to be resolved.

## Introduction

Between 50 and 40 thousand years ago (kya cal BP), the demographic landscape of Europe is transformed as Neandertals are replaced by anatomically modern humans (AMH) and disappear from the fossil record ^1^. Recent evidence from Bulgaria and the Czech Republic indicates that the first AMH arrived in Europe by at least 47-45 kya cal BP ^2–4^. At a continental scale, this would suggest a potential overlap of five to six thousand years between these human species ^3^. Yet, little is known about the nature, timing, and geographic areas of interaction between Neandertals and *Homo sapiens* during this critical period in human evolutionary history.

Archaeologically, the first part of this period – the Middle to Upper Palaeolithic transition – is characterised by so-called ‘Initial Upper Palaeolithic’ assemblages (e.g. Bacho Kiro, Temnata Dupka) and is increasingly interpreted as representing an initial, possibly unsuccessful migration of AMH into Europe occurring around 47-44 kya cal BP ^3,5,6^. The term ‘unsuccessful’ has been used as these initial groups appear to have left no visible genetic contributions to subsequent populations in Europe ^3,6^. The onset of the Aurignacian techno-complex *(sensu lato)* across Europe at around 42 kya cal BP is widely accepted as reflecting a second, more successful migration of AMH groups into Europe’s western extensions, and may signal the first *major* phase of European colonisation by our species ^5,7^. In many regions, Protoaurignacian and Early Aurignacian assemblages appear to rapidly replace so-called ‘transitional’ stone tool industries (e.g. Uluzzian, Châtelperronian, Lincombian-Ranisian-Jerzmanowician), some of which are considered to be products of Neandertals. Interestingly, genetic research indicates there to be significant variation in Neandertal ancestry for the earliest AMHs in Europe ^3,6,8,9^ and – although sample sizes are limited – it is revealing that no late European Neandertals have yet exhibited evidence of a recent modern human ancestor ^10^. One possible explanation for this pattern is that, at least in some regions, AMHs Europe may not have directly encountered Neandertals.

At present, the Châtelperronian stone tool industry of France and northern Spain shows the strongest association between a ‘transitional’ industry and Neandertal fossil remains. Despite the continued use of the ‘transitional’ moniker, however, it is now understood that this industry represents a fully ‘Upper Palaeolithic’ technological entity ^11–15^. Neandertal remains have been recovered from stratigraphic layers containing Châtelperronian artefacts at the two key French sites of Saint-Césaire and Grotte du Renne ^16–20^. However, the validity of these associations is debated, and consensus regarding both the makers of this industry and the reliability of the Neandertal associations is not unanimous ^11,21,22^. The other two French Neandertal specimens recovered from this period lack clear Châtelperronian associations (Les Cottés Z4-1514, La Ferrassie LF8), but have been directly-dated to between 43 and 40 kya cal BP ^10,23^. This is comfortably within the accepted chronological distribution of the Châtelperronian industry and overlaps substantially with the Saint Cesaire and Grotte du Renne Neandertals ^20,24^. Despite ongoing discussions, a Neandertal-attribution for Châtelperronian assemblages remains the most parsimonious and well-accepted model.

Technological similarities between some Châtelperronian and Protoaurignacian assemblages (i.e. blade and bladelet-based lithic technology, bone tools, and personal ornaments) ^13,14,18,25,26^ has led to discussion concerning the potential interactions between *Homo sapiens* and Neandertals in this region prior to the latter’s disappearance around 40 kya cal BP ^1,5,13,14,18,27–30^. Most notably, it has been proposed that the ‘Upper Palaeolithic’ character of Neanderthal Châtelperronian assemblages reflects the influence of allochthonous AMHs producing Protoaurignacian assemblages. However, whenever these two lithic industries are identified at the same site, Protoaurignacian assemblages are always located stratigraphically above Châtelperronian assemblages ^13^. In combination with chronological data suggesting an earlier ‘start’ date for the Châtelperronian, models which posit an Aurignacian influence as a causal mechanism for the *emergence* of the Châtelperronian appear to be presently unfounded. This does not, however, preclude the partial contemporaneity of these occupations *at some point in time.* In fact, Bayesian modelling of radiocarbon ages for Protoaurignacian and Châtelperronian assemblages in this region has already indicated that these occupations may have co-existed for upwards of 1600 years ^39^.

From a methodological perspective, two recent developments in the dating of archaeological phenomena are relevant to these discussions. The first concerns improvements to the calibration curve used to convert C14 measurements into reliable calendar dates ^32^. The recently operationalised IntCal20 radiocarbon calibration curve has significant implications for the chronology of the initial stages of the Upper Palaeolithic in Europe ^33^. Specifically, the identification of a radiocarbon time-dilation during the 48 to 40 kya cal BP time window – during which the radiocarbon clock appears to have run almost twice as fast as it should - has led to the suggestion that the European transition from Neandertals to AMH may have been a more compressed process which took place slightly earlier than previously thought **(Fig. 1)** (*ibid*.). This expanded C14 time scale was not accounted for in former calibration curves, and is thought to be related to a significant rise in atmospheric ^14^C production (on the order of as much as a 700% increase) linked to the transition into the Laschamp geomagnetic excursion, which reached its peak around 43-41 kya cal BP ^32^.

**Figure 1.**
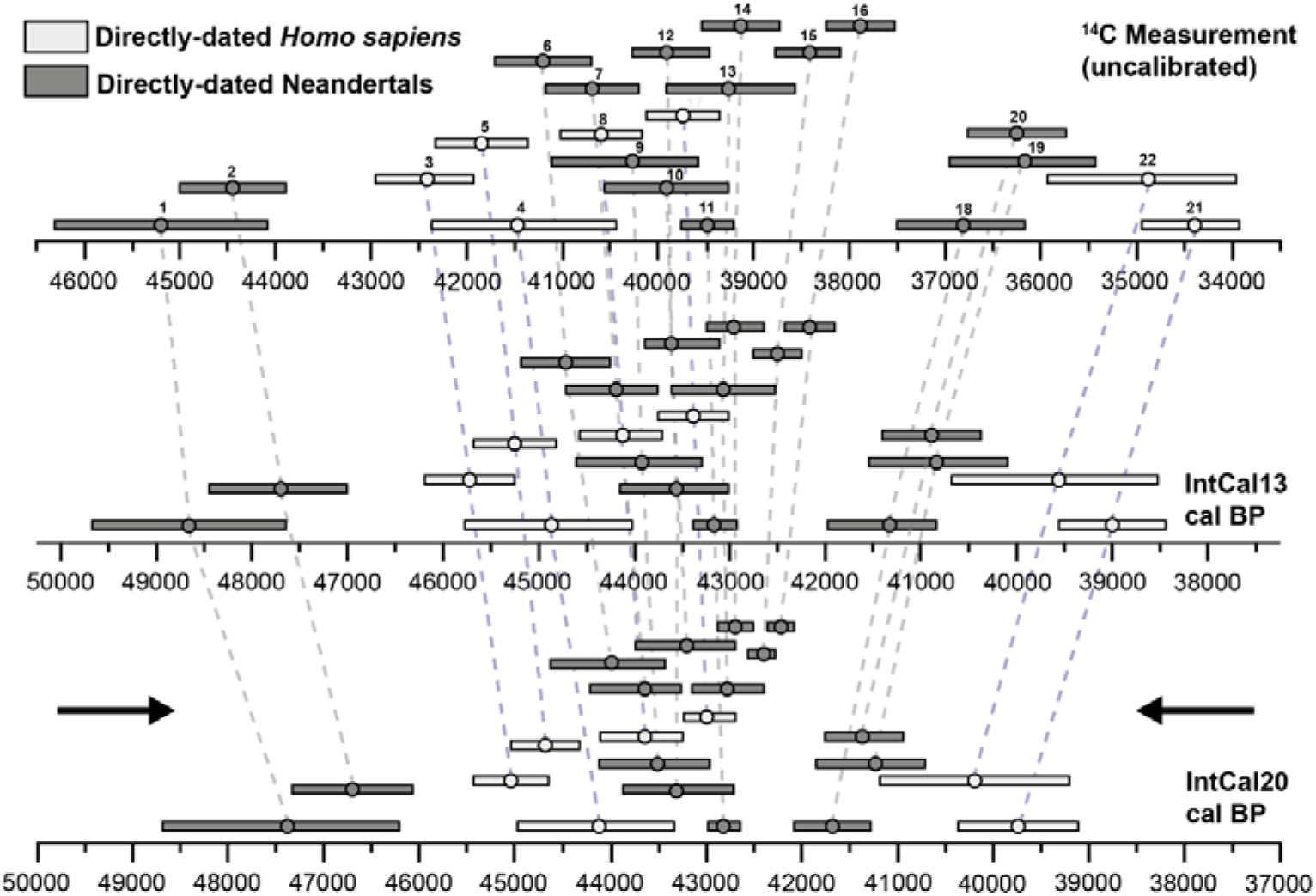
The effects of the recently operationalised IntCal20 radiocarbon calibration curve on C14 measurements produced for human remains between 50 and 37 kya (bottom) - compared with both the uncalibrated measurements (top) and the ages obtained using the previous generation curve (IntCal13, middle) (redrawn and adapted with permission after Bard et al., 2020). Note the ‘time-dilation’ causing a compression of dates centred around the 43-42 kya cal BP mark (black arrows). **1** – Les Rochers-de-Villeneuve (France), **2** – Vindija Cave Vi-33.26 (Croatia), **3** – Bacho Kiro (Bulgaria), **4** – Ust’-Ishim (Siberia), **5** – Bacho Kiro (Bulgaria), **6** – Goyet Q57-1 (Belgium), **7** – Goyet Q305-4 (Belgium), **8** – Bacho Kiro (Bulgaria), **9** – Neander Valley NN4 (Germany), **10** – Neander Valley Nean 1 (Germany), **11** – Les Cottés Z4-1514 (France), **12** – Goyet Q53-4 (Belgium), **13** – Neander Valley NN1 (Germany), **14** – Goyet Q376-1 (Belgium), **15** – Goyet Q56-1 (Belgium), **16** – Goyet Q55-1 (Belgium), **17** – Bacho Kiro (Bulgaria), **18** – Grotte du Renne AR-14 (France), **19** – Saint-Césaire (France), **20** – Spy 737a (Belgium), **21** – Tianyuan Cave (China), **22** – Pestera cu Oase (Romania). Figure was produced using Adobe Illustrator.

The second methodological development is the recent introduction of optimal linear estimation (OLE) modelling to archaeology from palaeontological and conservation sciences ^34^. OLE is a frequentist modelling approach that can reconstruct the full chronology of cultural and biological phenomena by statistically inferring origin (‘origination’) and end (‘extinction’) dates. Unlike traditional estimates which often use the earliest or latest known dated artefacts/fossils as a start or end point, OLE is able to infer how much longer a phenomenon is likely to have persisted prior to, or after, these known occurrences. In general terms, this method is underpinned by the assumption that we rarely, if ever, find the ‘first’ or ‘last’ occurrence of a species, artefact, or cultural tradition ^35,36^ Meaning that the earliest and latest instances of a given archaeological (or fossil) phenomenon are unlikely to ever be discovered and dated. OLE addresses this issue by using the temporal spacing of known artefact discoveries to statistically estimate the portion of the archaeological record that has not yet been, or is not able to be, discovered ^34,37^. In turn, providing a more accurate account of a phenomenon’s temporal presence.

These developments have potential to improve our understanding concerning the timing of the bio-cultural transformations characterising the Middle to Upper Palaeolithic transition, and western Europe serves as an ideal case study for their integration into discussions concerning the potential regional co-existence of Neandertals and AMHs. Of particular relevance is France and northern Spain, a region which features four Neandertal fossils directly-dated to between 44 and 40 kya cal BP ^10,20,23,24^, numerous well-studied and reliably dated Châtelperronian assemblages associated with late Neanderthals ^1,24,38–40^, and some of the earliest well-dated AMH-attributed Protoaurignacian contexts within western Europe ^41–45^.

Due to the sparsity of human fossil remains for this period, to address whether or not Neandertals and AMHs may have co-existed in any given region of Europe it is also necessary to evaluate whether the proxies used to define these groups in the archaeological record (assemblages, industries, techno-complexes etc.) can be considered geographically and temporally contemporaneous ^46^. Here, we use the recently operationalised IntCal20 calibration curve to recalibrate a large selection of C14 determinations for Châtelperronian assemblages, Protoaurignacian assemblages, and directly-dated late Neandertals from France, northern Spain, and Belgium. Then, we analyse these data using OLE modelling to statistically infer the appearance date of anatomically modern humans and the extinction date of Neandertals in this region. Finally, we compare the results of this approach with Bayesian models which rely on *known* dated occurrences as ‘start’ or ‘end’ points. By doing so, we provide a novel, testable hypothesis for the duration of overlap between Neandertals and *Homo sapiens* in this key region of western Europe.

## Results

The dataset consists of 56 uncalibrated radiocarbon age determinations from Châtelperronian and Protoaurignacian assemblages (n=28 and 28) from seven and ten archaeological sites, respectively. Collectively, covering northern Spain and south-west, central and Mediterranean France (**Supplementary Table S1, Supplementary Fig. S1**). In addition, to examine the temporal relationship of Neandertal fossils with these assemblages, we included all available radiocarbon estimations from directly-dated late (< 50 kya cal BP) Neandertal specimens within the surrounding region (France n=4, Belgium n=6, total n=10) (**Supplementary Table S1**). In total, 66 radiocarbon age determinations from 18 discrete, well-established archaeological sites are represented within the dataset (**Supplementary Table S1**). A detailed summary of the samples used here and the OxCal scripts used in the analysis, along with all accompanying information, is made available in full (**Supplementary Dataset S1, Supplementary Fig. S8**).

### Chrono-spatial patterning of known Châtelperronian, Protoaurignacian, and directly-dated Neandertal occurrences in the region

The plots summarising the distribution of the aggregated IntCal20 calibrated radiocarbon ages (at 95.4% confidence) for the Châtelperronian, Protoaurignacian, and directly-dated Neandertal datasets are illustrated in **Fig. 2** – including Bayesian start/end dates produced using the same datasets. The probability distributions show clear overlap between all three categories. Based on the aggregated datasets (**Supplementary Dataset S1, Supplementary Figs. S2-S4**), Bayesian modelling suggests a start date for to Châtelperronian between 45,343 and 44,248 kya cal BP, and an end date between 41,081 and 40,138 kya cal BP. The dataset for the regional Protoaurignacian produces a modelled start date between 42,873 and 41,747 kya cal BP, and an end date between 39,197 and 38,087 kya cal BP. For the directly-dated Neandertal dataset, the modelled end date for Neandertal presence in this region is predicted to have occurred between 41,757 and 39,859 kya cal BP. Taken together, the chronological data for the regional Protoaurignacian, Châtelperronian, and directly-dated Neandertals show a partial overlap. For example, calibrated age ranges produced for the Protoaurignacian assemblages at Isturitz (n=4), Labeko Koba (n=2), Gatzarria (n=1), Esquicho-Grapaou (n=1), and L’Arbreda (n=4) overlap either entirely or near-entirely with three directly-dated Neandertals from France – Saint-Césaire (42206-39960 cal BP, IntCal20), La Ferrassie (LF8, 41696-40827 cal BP, IntCal20), and Grotte du Renne (AR-14, 42370-40778, IntCal20).

**Figure 2.**
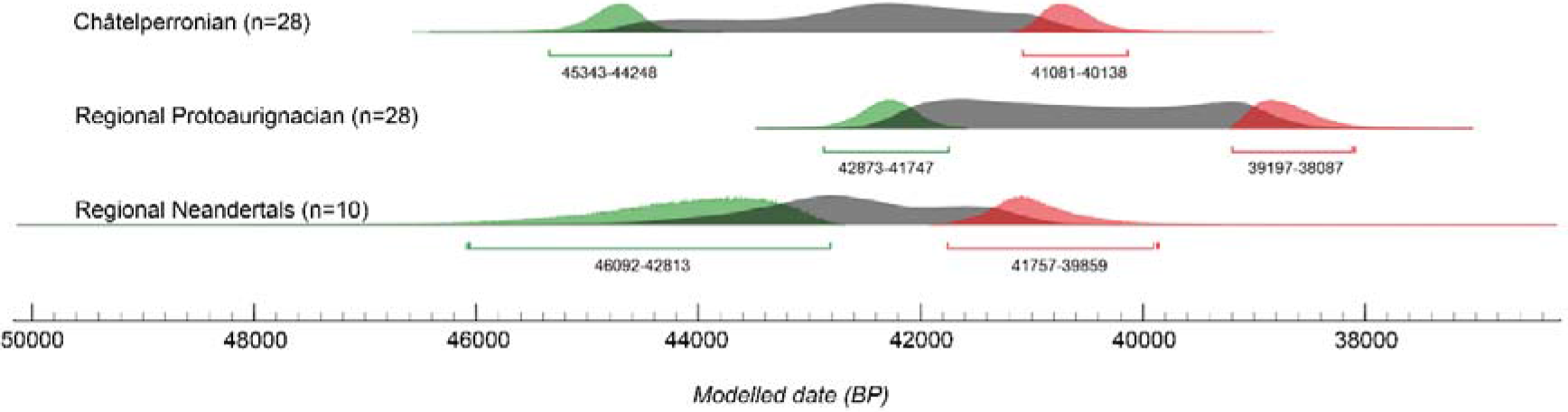
Kernel Density Estimation plots and Bayesian start/end date probabilities summarising the distribution of the aggregated calibrated radiocarbon ages for a) the Châtelperronian assemblages (n=28), b) the Protoaurignacian assemblages (n=28), and c) the directly-dated late Neandertals (n=10) included in this study. Bayesian start and end dates are visualised in green and red, respectively. Figure was produced in the OrAU OxCal software (Ramsey, 1995, v4.4).

In terms of which sites are accounting for this overlap, for the Protoaurignacian sites the calibrated age ranges with the oldest potential ages derive from: Isturitz (OxA-X-2694-17, OxA-23435, OxA-23436, OxA-23434), Labeko Koba (OxA-21766, OxA-X-2314-43), Gatzarria (OxA-22554), Esquicho-Grapaou (OxA-21716), and L’Arbreda (OxA-21784, OxA-21665, OxA-21664) (**Supplementary Fig. S4**) – forming a coherent geographic cluster at the southern limit of the study region (**Fig. 3, a-f**). This pattern suggests that the early stages of the first modern human settlement of this region likely followed a south-north pattern of occupation – with the Protoaurignacian progressively appearing further north and replacing the Châtelperronian in stratigraphic sequences (**Fig. 3, d-f**).

**Figure 3.**
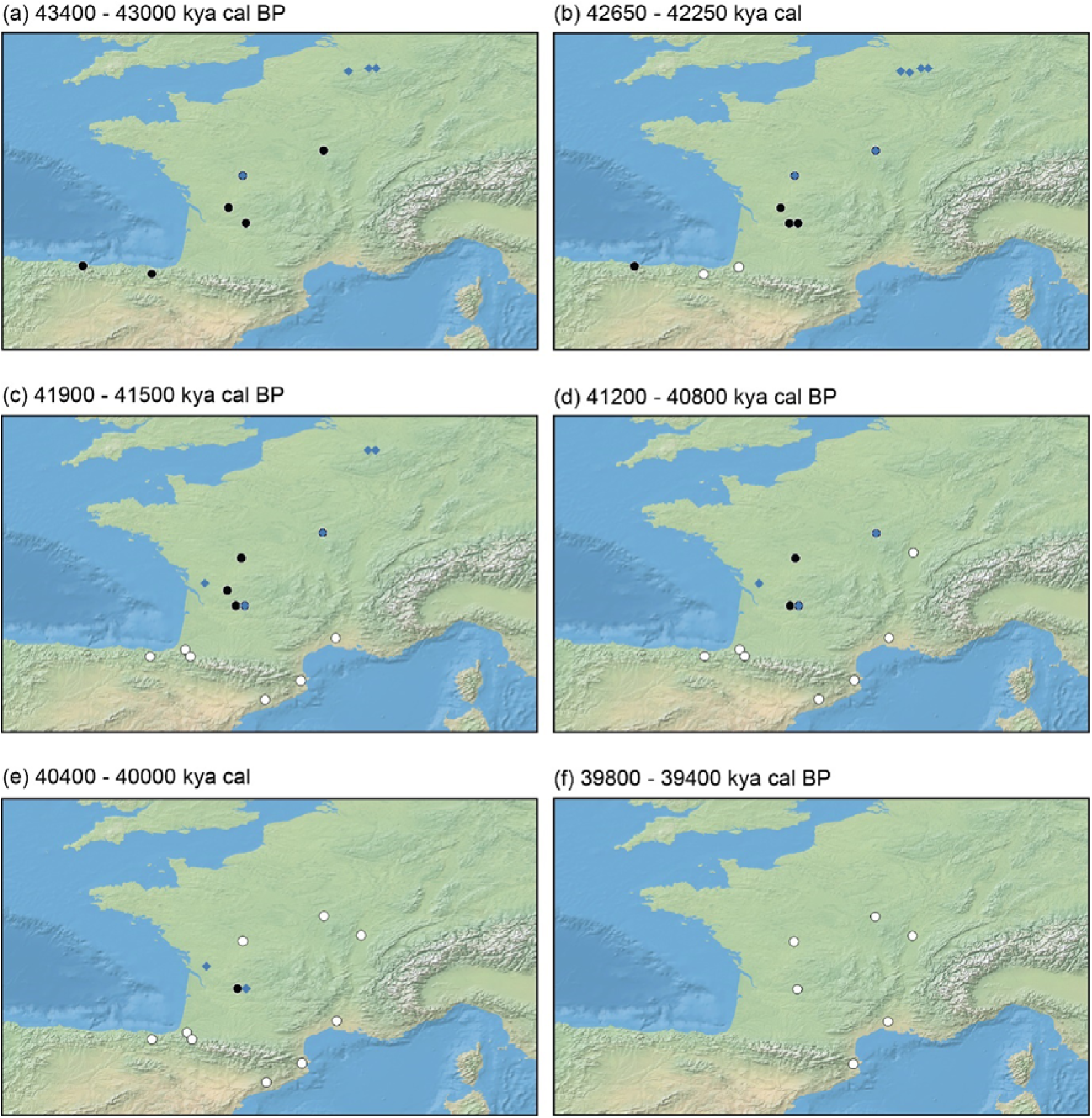
Geographic appearance of dated occurrences for the Châtelperronian (black circles), Protoaurignacian (white circles), and directly-dated Neandertals (blue diamonds) in the study region between 43,400 (a) and 39,400 (f) years cal BP. Figure was produced using the ‘spatio-temporal modeller’ function in the QGIS software (v4.4) (QGIS Geographic Information System, QGIS Association, 2021) and compiled in Adobe Illustrator.

### Using OLE modelling to infer the ‘emergence’ time of the Protoaurignacian and the ‘extinction’ time of the Châtelperronian and Neandertals in the region

We had three objectives, with each requiring its own OLE model and respective sample (see ‘Methods’):

1. *Estimating the emergence date of the Protoaurignacian in France and northern Spain.* The nine oldest Protoaurignacian dates from nine discrete archaeological sites are entered into this OLE model (**Supplementary Table S2, Supplementary Fig. S7**) which is run in the reverse temporal direction.
2. *Estimating the end date of the Châtelperronian in France and northern Spain.* The eight youngest Châtelperronian dates from seven discrete archaeological sites are entered into this OLE model (**Supplementary Table S3, Supplementary Fig. S6**) which is run in the forward temporal direction.
3. *Estimating the extinction date of regional Neandertals.* Ten direct dates of late Neandertal individuals from France (n=4) and Belgium (n=6) are entered into this OLE model which is run in the forward temporal direction (**Supplementary Table S4, Supplementary Fig. S5**).

OLE modelling infers the Protoaurignacian to have likely emerged in France and northern Spain by 42,269 to 42,653 years cal BP. The upper bound of this T_O_ date range is defined by the resampling technique, while the lower uses the central tendency (mean) dates derived from the C14 date range. As explained earlier, we consider the resampling estimates to better account for the range uncertainly inherent to C14 dating. T_CI_ dates, beyond which the Protoaurignacian only has a 5% chance of preceding this point, provide a bracket of 43,394 – 44,172 years cal BP. Upper and lower bounds were again defined by the resampling technique and central tendency dates (respectively). OLE modelling estimates the Châtelperronian to have disappeared by 39,894 to 39,798 years cal BP. The upper bound of this T_O_ date range is defined by the resampling technique, while the lower uses the central tendency (mean) dates. T_CI_ dates, beyond which the Châtelperronian only has a 5% chance of following this point, provide a bracket of 37,838 - 37,572 years cal BP. Again, upper and lower bounds were defined by the resampling and central tendency dates respectively. OLE modelling infers the localised extinction of Neandertals in France and Belgium to have occurred between 40,870 to 40,457 years cal BP. The upper bound of this T_E_ date range is defined by the resampling technique, while the lower uses the central tendency (mean) dates. T_CI_ dates, beyond which Neandertals only have a 5% chance of following this point, provide a bracket of 39,688 to 38,752 years cal BP - with the upper and lower bounds again defined by the resampling and central tendency techniques, respectively. Across all OLE models, the resampling approach extended temporal ranges by several hundred years compared to the central tendency (mean) based estimates. The results of the 10,000 resampling iterations for each model are illustrated in **Fig. 4** and the raw data is available in full (**Supplementary Data S2**). Combined, OLE modelling suggests the Protoaurignacian to have emerged around 1399 – 2196 and 2375 – 2855 years before Neandertals and the Châtelperronian industry (respectively) disappeared from the region.

**Figure 4.**
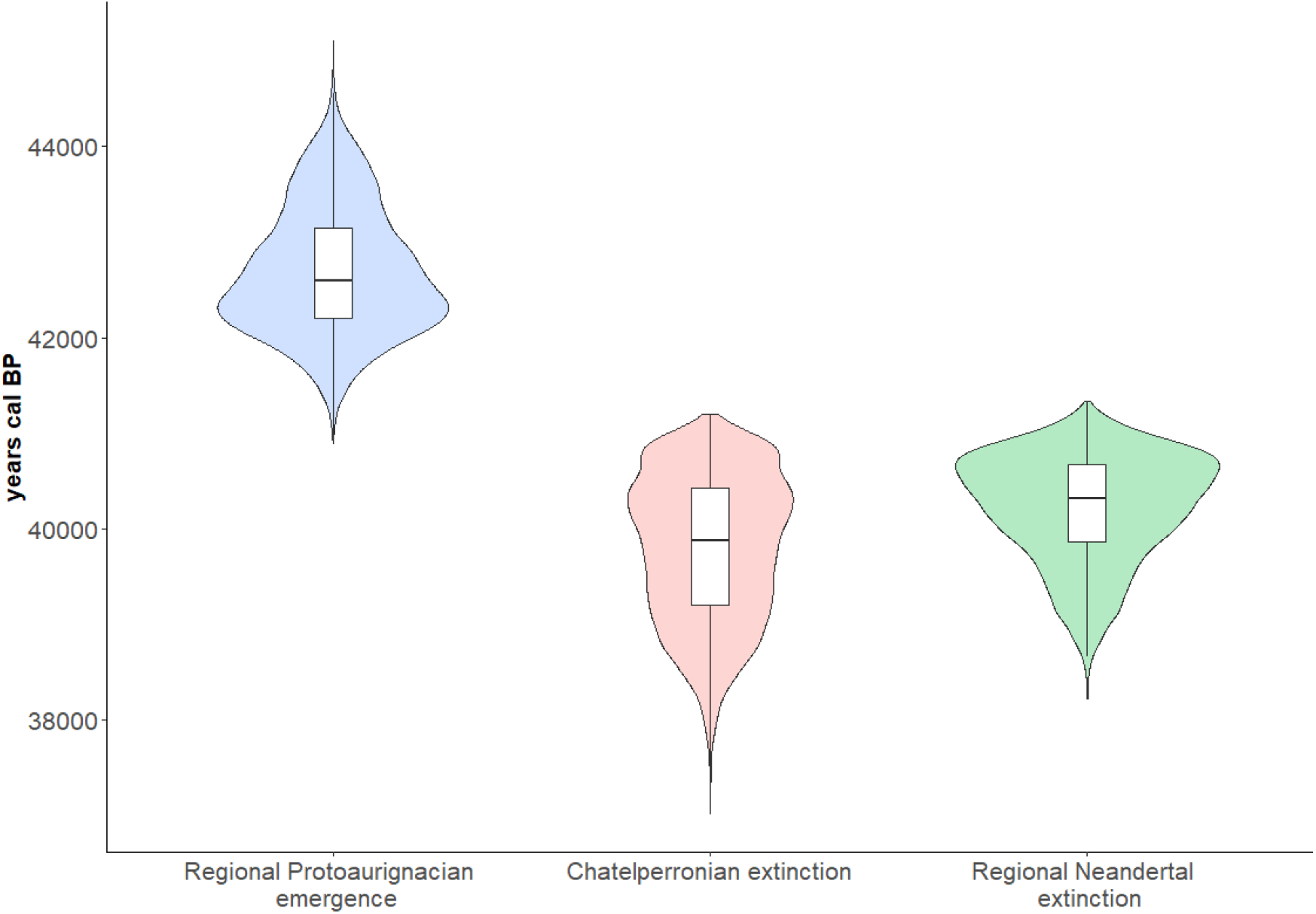
Resampling results for the three OLE models: modelled Protoaurignacian regional emergence time (left), modelled Châtelperronian extinction time (centre), and modelled Neandertal regional extinction time (right). The horizontal bar in each respective boxplot represents the mean value of the 10,000 resampling iterations referred to in the text. Figure was produced using the ‘ggplot2’ package in R (version 4.0.3).

## Discussion

Based on OLE modelling of their respective ‘origination’ and ‘extinction’ dates, the Protoaurignacian potentially appeared around 1400 – 2900 years before Neandertals and the Châtelperronian industry disappeared from France and northern Spain. These results raise the possibility of an extended period of co-existence between AMH and Neandertals in this region. Additionally, and as has been previously suggested, the chronological overlap between these occurrences appears to be geographically structured. The oldest calibrated age ranges from well-dated Protoaurignacian assemblages initially form a cluster at the southern and northern limits of France and Spain respectively, overlapping with dates produced for Châtelperronian assemblages in the central-northern parts of France. This suggests that the chronological overlap may have been geographically structured, with the Protoaurignacian following a south to north pattern of appearance. Moreover, based on OLE estimates produced using directly-dated Neandertal fossil remains, the onset of the regional Protoaurignacian is modelled to have preceded the extinction of Neandertals in this region by upwards of 2200 years.

These results are perhaps not surprising given the nature of probability ranges for calibrated radiocarbon determinations produced for this period – which is temporally situated near the upper acceptable limit of radiocarbon dating (circa 50 kya) ^47^. However, the fact that the OLE ‘extinction’ and ‘emergence’ estimates produced here do not go far beyond the ranges identified in the calibrated radiocarbon dates themselves is notable, and is directly related to the temporal spacings observed for each of the occurrences. In each case the latest/oldest series of dates for each category (Protoaurignacian, Châtelperronian, Neandertal) reflect a narrow temporal band with little variation and inter-date spacing (ie., the dates are chronologically close), after which no additional dated occurrences are known. This has two potential implications. Firstly, that the oldest and/or youngest dates for each industry are *likely close* to the true emergence and/or extinction date of that industry ^34^. Secondly, and relatedly, the true emergence and/or extinction dates may in some cases be slightly *more conservative* than the upper limits of the oldest and/or youngest calibrated dates themselves. This is perhaps particularly relevant for the Châtelperronian, which is widely acknowledged as largely reflecting relatively ephemeral and short-lived occupations ^13^ – with the exception of some notable examples ^14,24^.

Of course, there are limitations in this analysis which require consideration. The most obvious concerns the sample size of archaeological sites included in this work – which was dictated by the decision to employ strict, conservative sampling requirements for the radiocarbon datasets. And while we acknowledge that the sample considered here reflects only a portion of known Châtelperronian and Protoaurignacian occurrences within this region, it does cover their *known geographic distribution.* Moreover, OLE works best with limited datasets, such as this. A second potential limitation concerns the radiocarbon determinations themselves. Any model is, of course, only as reliable as the data entered into it. The assumption taken here is that the age ranges entered into the models reflect meaningful datapoints for the chronological presence of these occurrences. This, in time, may change as the duration of these industries is increasingly refined. At present however there is no clear evidence to doubt the reliability of the radiocarbon determinations used in this study, but future work may necessitate the revision of this model as more sites are dated - or re-dated - and further methodological advancements are made.

From an archaeological perspective, of relevance to these results is the acknowledged presence of bladelet technologies, osseus artefacts, and personal ornaments within a growing number of Châtelperronian and Protoaurignacian contexts. Unanimously seen as a trademark of the Protoaurignacian techno-complex (with the laterally retouched Dufour bladelet [sub-type Dufour] *fossile directeur* commonly constituting a substantial portion of Protoaurignacian assemblages), evidence for some form of intentional bladelet production and/or modification within the Châtelperronian has now been reported from at least four open-air sites ^12,15,48^ and six cave sites ^14,40,48–51^. To what extent (if any) these similarities represent some form of connection between these industries remains unclear, but the potential contemporaneity of the groups producing these assemblages is certainly of relevance.

Of course, the results presented here do not aid in answering the question of which human group(s) were responsible for producing these industries, but the temporal and geographic proximity of directly-dated Neandertal remains to both Châtelperronian and AMH-attributed Protoaurignacian assemblages in the region is – in the current state of knowledge – difficult to overlook. With this being said, the recent chronological re-evaluation of late-dating Belgian Neandertals has convincingly demonstrated that they are likely substantially older than previously thought **(Fig. 5)**^52^. With this development, the Neandertals from France included in this study are now among the latest directly-dated Neandertals identified throughout the inferred geographic distribution of this human group. This raises an important consideration: it is possible that future work employing emerging radiocarbon dating techniques designed to further mitigate anthropogenic and/or natural contamination issues (e.g. Compound Specific Radiocarbon Analysis) may, in time, either confirm or revise their currently accepted ages.

**Figure 5.**
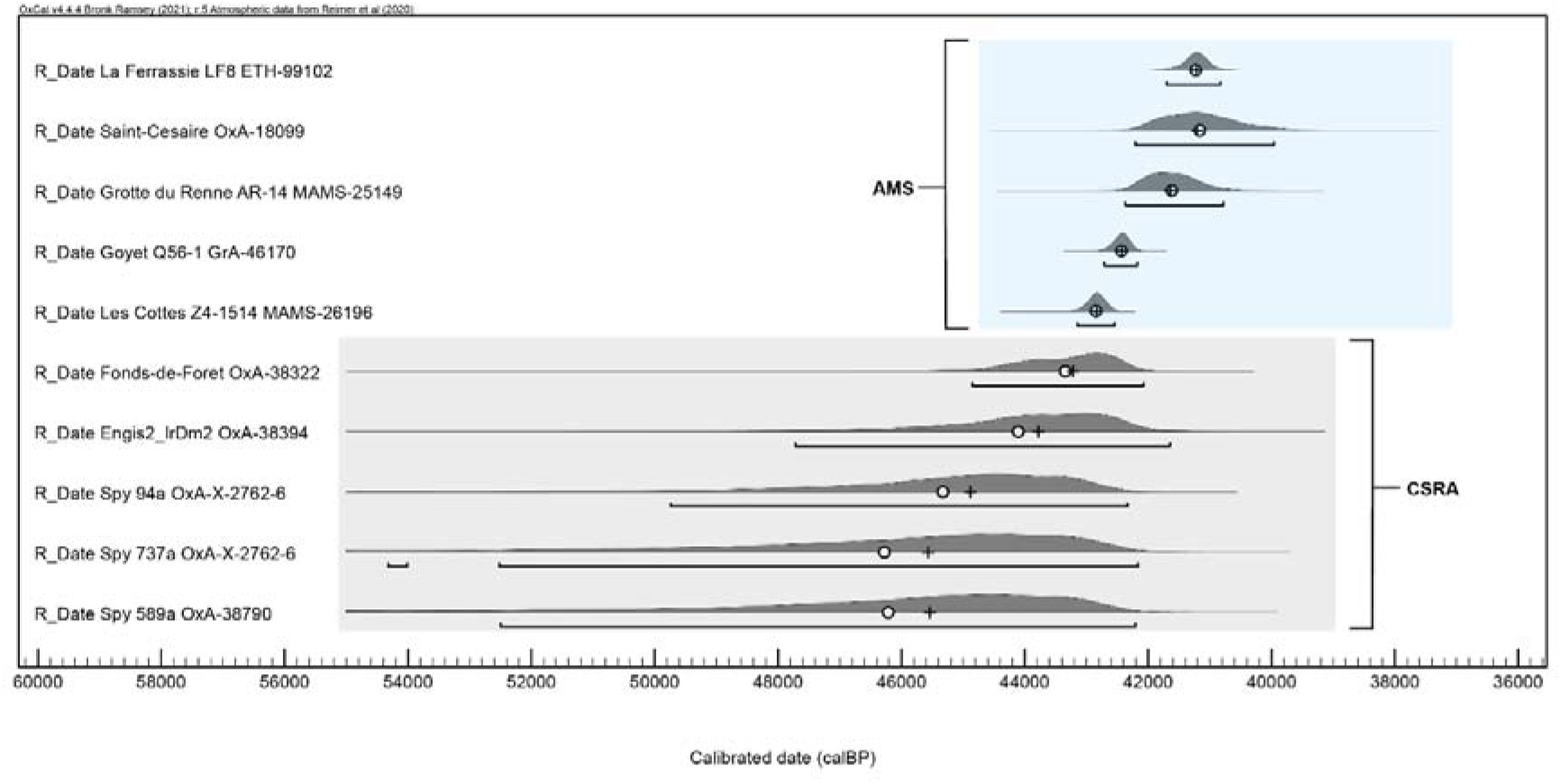
Calibrated age ranges for the ten late Neandertals included in this study. The lower five specimens were dated using compound specific radiocarbon analysis (CSRA) of hydroxyproline (Deviese et al., 2021) while the upper five specimens were dated with the AMS method. All samples were prepared using ultrafiltration. Figure was produced in the OrAU OxCal software (Ramsey, 1995, v4.4) and compiled using Adobe Illustrator.

Regardless, considering the rapidly evolving understanding of the European demographic landscape preceding the onset of the Aurignacian techno-complex *sensu lato* ^3,4,6,8^, it is clear that more work is needed to evaluate the biological identity and the cultural connections, if any, between the makers of archaeological industries across the European landmass during this period. However, at present, the only hominin species to as of yet be securely associated with Châtelperronian assemblages, based on both morphological and genetic evidence, is Neandertals. For the Protoaurignacian, the case is reversed – with the only published hominin association in a Protoaurignacian context being two deciduous teeth from Riparo Bombrini and Grotta di Fumane caves in Italy which have been attributed as *Homo sapiens* based on morphological criteria and mitochondrial DNA, respectively ^53^. With this said, at present, the reality is that most Protoaurignacian assemblages are serving simply as *well-accepted proxies* for the presence of *Homo sapiens* – but the validity of this unilateral association is, in the current state of evidence, far from certain. In fact, in many ways the same can be said for Châtelperronian assemblages and their unilateral association with Neandertals ^11,22,51^. With specific reference to the ongoing proliferation of paleogenetic research (including sedimentary aDNA analysis) and the enormous increase in efficiency for Zooarchaeology by Mass Spectrometry (ZooMS), future work will undoubtedly shed new light on the biological makers of these industries.

In a similar vein, the ‘origins’ of the Châtelperronian industry remains an open question – but it is becoming increasingly evident that models which posit an Aurignacian influence as a *causal mechanism* for the *emergence* of the Châtelperronian are chronologically and stratigraphically unfounded. The onset of the Châtelperronian, in the current state of knowledge, appears to clearly pre-date the appearance of the Protoaurignacian - both regionally and at a European scale. However, the spatio-temporal overlap of these assemblages in France and northern Spain – and their potential overlap with multiple directly-dated Neandertals from the region - lend credence to the idea that the early stages of the Upper Palaeolithic in this region may have involved the proximal co-existence of different human groups, likely irrespective of their biological classification.

## Conclusion

Optimal linear estimation modelling predicts the appearance of *Homo sapiens* and the Protoaurignacian in France and northern Spain by 42,269 to 42,653 years cal BP, and the ‘extinction’ of the Châtelperronian and regional Neandertals by 39,894 to 39,798 and 40,870 to 40,457 respectively – suggesting a possible overlap of around 1400 to 2800 years between these human groups in the region. In addition, this chronological overlap appears to be geographically structured, with the Protoaurignacian following a south to north pattern of appearance. Taken together, these observations strengthen the proposition that the initial Upper Palaeolithic in this region likely involved the extended co-existence of Neandertals and *Homo sapiens.* the precise nature of this co-existence, however, remains to be resolved.

## Methods

### Site and sample selection

The lack of adequate pre-treatment procedures for many of the radiocarbon age determinations produced prior to the 2000s has led some to suggest that many, if not all, of these early dates should be considered unreliable ^5^. As a result, and in line with this proposition, we took a conservative approach to the site and sample selection for this study. Only modern (year 2000 onwards) radiocarbon dates produced on a) anthropogenically modified or unmodified bone samples, b) tooth samples, and c) antler samples were included. In addition, all samples were prepared using the ultrafiltration pre-treatment protocol ^54–59^ and all age determinations except for five directly-dated Belgian Neandertals – dated using compound specific radiocarbon analysis (CSRA) of hydroxyproline (HYP) ^52^ – were produced using the AMS radiocarbon dating method ^56^. To further ensure the quality of the dataset, all samples included here have reported and fulfilled the requirements of well-accepted collagen quality control measures (C:N ratios, %C, %N, % of collagen, d13C, and d15N) considered necessary to establish the lack of contamination and/or degradation of collagen ^60,61^.

### IntCal20 calibration, chronological distribution summaries, and chrono-spatial patterns

All 66 radiocarbon age determinations were calibrated in the OrAU OxCal software ^62^ using the IntCal20 calibration curve ^32,63^ to produce age ranges in calendar years before present (BP) at 95.4% confidence. We used Kernel Density Estimation (KDE) in combination with Bayesian start/end date modelling – both included within the OxCal software (v4.4) ^64^ – to summarise the distributions of each occurrence based on the available chronological data. Of course, radiocarbon age determinations retain a degree of uncertainty that is expressed by a radiocarbon-date distribution. Bronk Ramsey^64^ has proposed an algorithm to incorporate that uncertainty into a KDE. This algorithm samples the individual radiocarbon age ranges to produce a set of probable dates, one for each event in a given database (calibrated age range). The algorithm then applies a KDE to the random sample of dates to produce a smooth estimate of temporal event density. This process is repeated for ten thousand iterations and the resulting average constitutes the final KDE model. While this method produces an accurate summary of available chronological information, it does not *necessarily* provide a true representation of through-time variation in occurrence-counts. This is to say, high and low points in the density distribution do not necessarily reflect a true increase or decrease in the through-time presence of the occurrence in reality, as radiocarbon datasets always represent an incomplete sample of a phenomenon. KDE and single phase Bayesian start/end date modelling were used to summarise and compare the distribution of calendar age probability ranges for all Châtelperronian and Protoaurignacian assemblages (n = 28 dates each), along with the 10 directly-dated late Neandertals. This approach synthesises and compares the aggregated chronological data for each occurrence, and does not seek to establish multi- or single-phase Bayesian models for any given site. Instead, it seeks to a) evaluate the general temporal trends within the chronological datasets for each category (Protoaurignacian, Châtelperronian, directly-dated Neandertals), b) identify the degree of overlap between these occurrences based on their aggregated datasets, and c) frame the results of this more traditional approach with those of the OLE modelling. Both the scripts used for this analysis and their output is available in the Supplementary Information (**Supplementary Figs. S2-S4, S8**).

To examine any geographic patterning within the chronological data, we created time-slice visualisations using the inbuilt ‘spatio-temporal modeller’ function in the QGIS software (v4.4) (QGIS Geographic Information System, QGIS Association, 2021). The dataset used for this visualisation consists of all IntCal20 calibrated radiocarbon age determinations (at 95.4% confidence) produced during the preceding step (66 dates from 17 discrete archaeological sites). The maximum possible age range of each occurrence within a given site (i.e. Châtelperronian, Protoaurignacian, directly-dated Neandertal) is used as the unit of analysis. In other words, the oldest and youngest possible date for each occurrence at a site act as the chronological boundaries for its presence at that site. As a result, these boundaries should not be taken as reflecting ‘true’ occupational durations at any given site. The intention of this approach is not to propose occupational durations, but to a) identify the geographic regions in which the earliest dates for Protoaurignacian assemblages appear to occur and b) highlight where any chronological overlap between the Châtelperronian, Protoaurignacian, and direct Neandertal age determinations appears to be manifested geographically.

### Inferring ‘origination’ and ‘extinction’ dates using optimal linear estimation modelling

OLE uses the timing and chronological spacing of known archaeological occurrences to statistically estimate how much earlier or longer the phenomenon is likely to have existed beyond the current known archaeological record (i.e. beyond known dated sites). It requires the oldest or youngest currently known dated occurrences of a phenomenon to be entered into the model, depending on whether it is being used to estimate an ‘origin’ or ‘end’ date. Estimated ‘origin’ and ‘end’ dates rely on the assumption that the dates entered into the model display (at least roughly) a joint distribution with a ‘Weibull form’. The form (shape parameters) of the Weibull distribution in the OLE model is estimated based on the chronology (spacing) of the dates entered into the model. From which an ‘end’ or ‘origin’ point can be produced, depending on the temporal direction of the model. Ten dates are generally considered as optimal for OLE ^37,65,66^, although it has been applied to lower sample sizes, with datasets of five having demonstrated good accuracy ^1^.

It is important to note that although first developed for conservation science ^37,65^, OLE has no parameters specific to biological organisms and can be readily applied to cultural traditions ^67^. The robusticity of OLE has been repeatedly demonstrated within a variety of scenarios, including those that vary in temporal scale, ‘sighting’ probabilities, and search efforts and trajectories ^66,68^. This means that OLE is likely reasonably accurate in providing ‘origin’ and ‘end’ estimates in most archaeological scenarios. As with any statistical modelling, results are only as accurate as the data entered into the model, and if there is uncertainty in the archaeological records used then this will be reflected in the security of the estimated dates. However, if there are securely dated sites to sample and all of the model’s assumptions are met ^34^, then “generally precise and accurate estimates” can be assumed ^68^.

OLE modelling is particularly amenable to dating archaeological phenomena as it works well with sparse datasets and only needs to consider the most recent or earliest records of a cultural tradition. In other words, the accuracy of the model’s result is not increased through the inclusion of large numbers of dated sites. As such, the datasets used for the models presented here consist of the youngest and oldest (depending on the direction of the model) calibrated age ranges of a particular archaeological occurrence. Each data-point included in an OLE model should represent a discrete occurrence of the phenomena in question. Therefore, each cultural occurrence (stratigraphic layer) was only represented by one datapoint (calibrated age range) as it is generally impossible to tell whether other dates produced within the same context can be considered as representing a discrete occurrence of that phenomenon. When overlap did occur, preference was given to the oldest or youngest dated sample, depending on the direction of the model. This is a relatively conservative way of defining discrete occurrences within OLE modelling ^34^. Our objectives here were to use OLE modelling to estimate the ‘origin’ date of the Protoaurignacian in France and northern Spain, the ‘extinction’ date of the Châtelperronian in the same region, and the ‘extinction’ date of Neandertals in the local and surrounding region. The three objectives and their required datasets are summarised here:

#### Estimating the start date of the Protoaurignacian in France and northern Spain

This model requires the *oldest* Protoaurignacian dates from the region, with the model run in the reverse temporal direction. The oldest available date for each well-dated Protoaurignacian site is used as a unit of analysis. We chose to exclude the oldest radiocarbon determination from the Protoaurignacian at Isturitz (OxA-X-2694-17) due to its low collagen yield (<1mg) and unclear depositional history ^42^. Additionally, we also chose to exclude two dates from Trou de la Mère Clochette (OxA-19622 and OxA-19621) produced on fragments of split-based points (antler) due to the uncertainty of their proposed cultural designation to the Protoaurignacian ^69^.

#### Estimating the end date of the Châtelperronian in France and northern Spain

This model requires the *youngest* Châtelperronian dates from the region, with the model run in the forward temporal direction. The youngest available date for each Châtelperronian context is used as a unit of analysis. We chose to include two radiocarbon dates from Grotte du Renne, as this site preserves multiple Châtelperronian layers ^24^.

#### Estimating the localised extinction date of late Neandertals

This model requires all reliably-produced direct dates of late Neandertal individuals from the broader region (France n=4, Belgium n=6) to be entered into an OLE model run in the forward temporal direction.

To account for C14 dating producing date ranges with even likelihood, and in line with research that has shown that mean values are an unreliable approach for summarising calibrated radiocarbon age ranges ^70^, we apply a resampling approach to the OLE modelling in which individual dates are randomly drawn, with uniform distribution, between the upper and lower age limits for each calibrated age range. These randomly generated datasets are in turn entered into the OLE model, and this process is repeated for 10,000 iterations ^67^. The mean value from these 10,000 iterations is then used as the origin or end date for this resampling approach. Given the large uncertainties produced for calibrated radiocarbon dates belonging to this period, we consider this approach as being both more statistically robust and inferentially cautious than the alternative (central estimate technique) approach, which utilises the mean date of each calibrated age range as a unit of analysis (i.e. the resampling approach does not depend on a single [mean] value as a datapoint for a calibrated age range which often spans multiple thousands of years) ^34^.

The model’s formulaic expression is available in the original articles describing OLE ^37,65^, along with more recent open access archaeological articles ^67,71^. All models were run in R (version 4.0.3) using the sExtinct software package ^68^. For the ‘origin’ dates the models were adjusted to run in the reverse temporal direction to those provided by Clements ^68^. The 10^th^ youngest or oldest dates were used as the beginning of the period, dependent on the direction of the model. Two estimated dates were produced by each model. One represents the estimated origin (T_O_) or end (T_E_) date of the phenomenon in question. The other represents the upper bound of each model’s confidence interval (*T*_CI_). *T*_O_ and *T*_E_ dates are the main output of the OLE models and are represented here as years before present (BP). *T*_CI_ dates represent the point beyond which the probability of the phenomena existing prior to or after this point in time, depending on the direction of the model, has a 5% or less probability (i.e., α□=□0.05).

## Acknowledgements

We would like to give a warm thank you to everyone within the Human Origins research group at Leiden University for their valuable feedback and stimulating conversations. We also thank Edouard Bard for his permission to reproduce a figure used in this text. This research is funded by the Dutch Research council (NOW) ‘Neandertal Legacy’ grant (VI.C.191.070) awarded to M. Soressi. This paper was presented as a Pecha Kucha during the annual ESHE meeting in 2021.

## Author contributions

ID, AK, and MS conceptualised the study. ID performed all data collection, Bayesian analyses, and GIS analyses. AK ran the OLE models and contributed text. ID wrote the main manuscript text and prepared all figures. ID, AK, and MS reviewed the manuscript.

## Data availability statement

All data analysed and generated during this study are included in this published article (and its Supplementary Information files).

## Additional information

The authors declare no competing interests.

