## Supplementary Online Material for "Optimal linear estimation models predict 1400-2800 years of co-existence between Neandertals and *Homo sapiens* in western Europe"

**Supplementary Figure S1.** The archaeological sites in France and northern Spain considered in this study. **1** - La Güelga, **2** - Labeko Koba, **3** - Isturitz, **4** - Gatzarria, **5** – Abric Romaní, **6** – L’Arbreda, **7** – Esquicho-Grapaou, **8** – La Ferrassie, **9** – Cassenade, **10** – La Quina-Aval, **11** – Saint-Césaire, **12** – Les Cottés, **13** – Grotte du Renne, **14** – Trou de la Mère Clochette, **15** – Spy, **16** – Goyet, **17** – Engis, **18** – Fonds-de-Forêt.

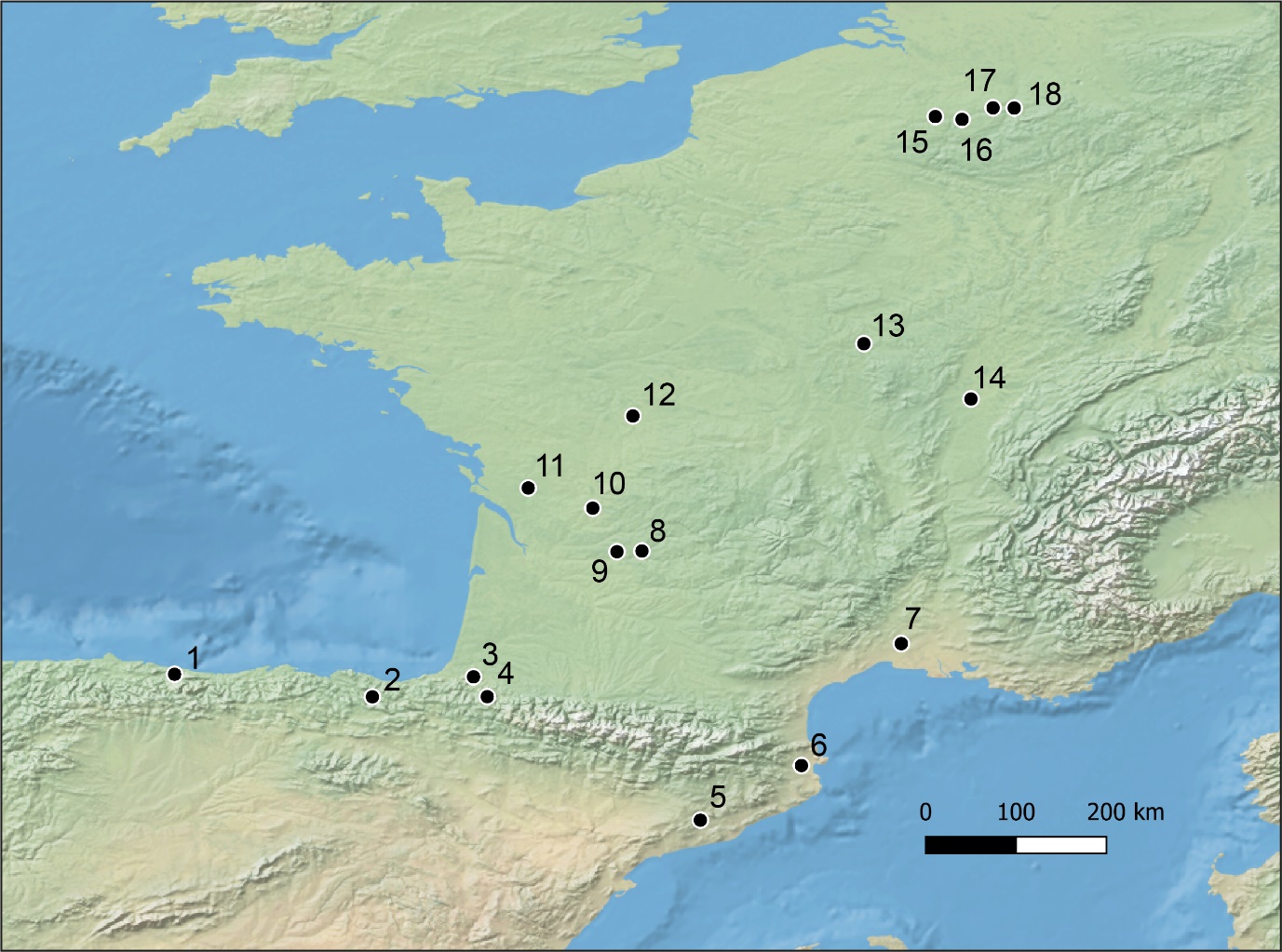

**Supplementary Figure S2.** Modelled Bayesian end dates for the total sample of directly-dated Neandertal radiocarbon determinations (n=10) used in this study, and produced in OxCal (v4.4) (Bronk Ramsey, 2006).

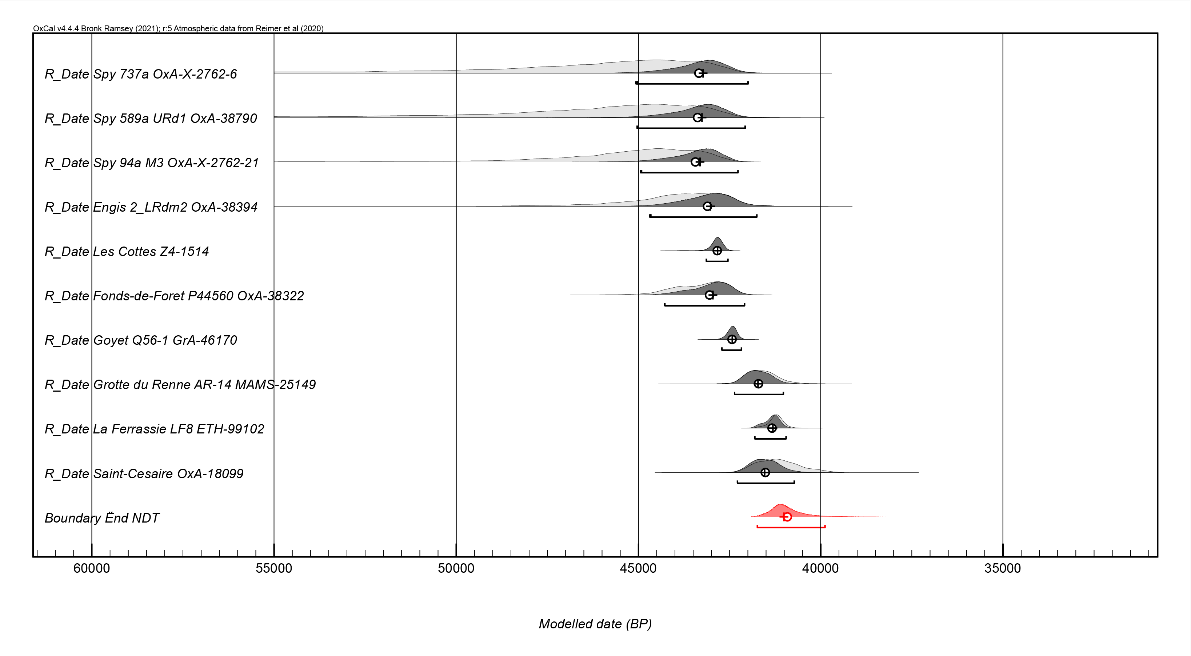

**Supplementary Figure S3.** Modelled Bayesian start and end dates produced in OxCal (v4.4) (Bronk Ramsey, 2006) for the Protoaurignacian radiocarbon dataset (n=28).

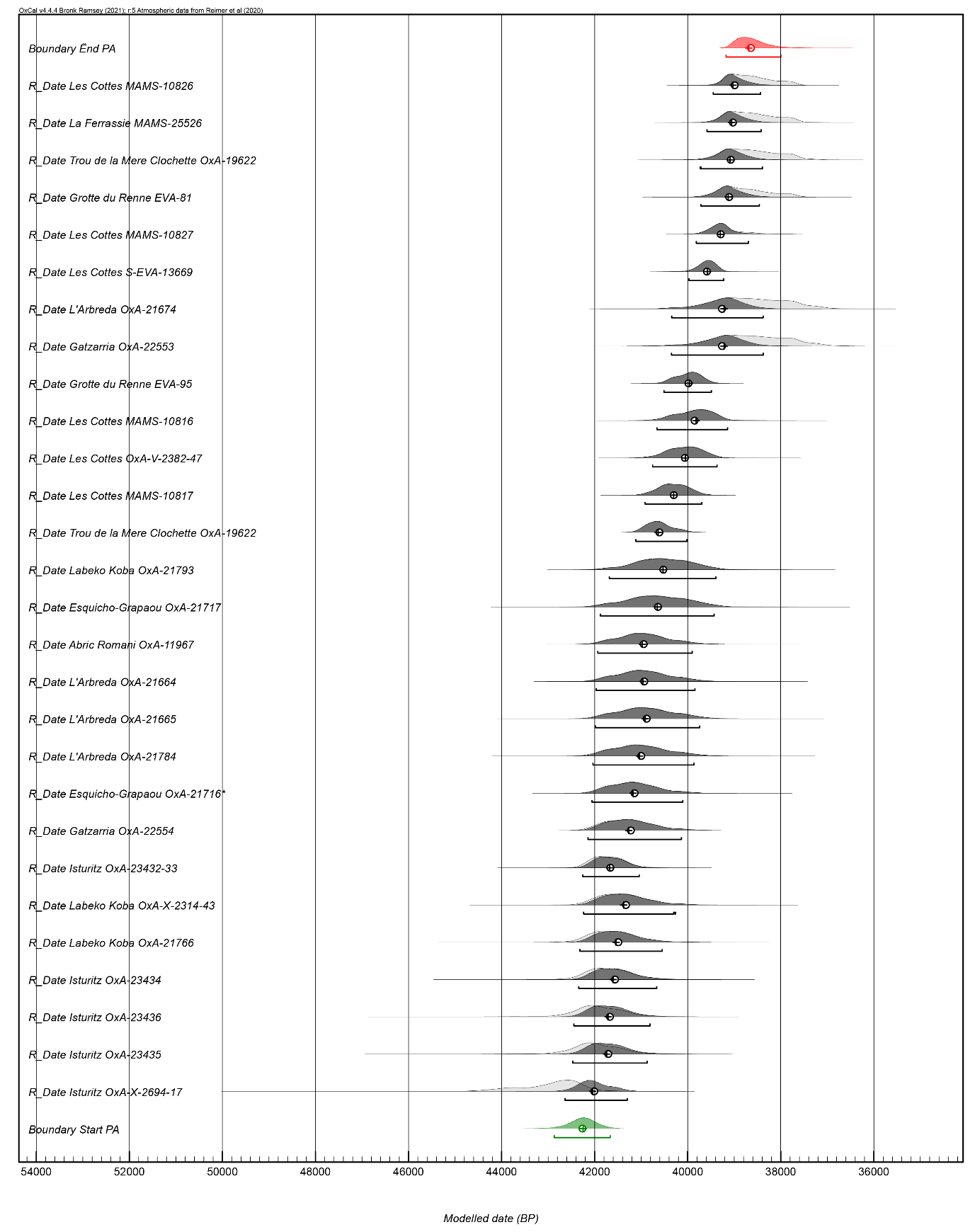

**Supplementary Figure S4.** Modelled Bayesian start and end dates produced in OxCal (v4.4) (Bronk Ramsey, 2006) for the Châtelperronian radiocarbon dataset (n=28).

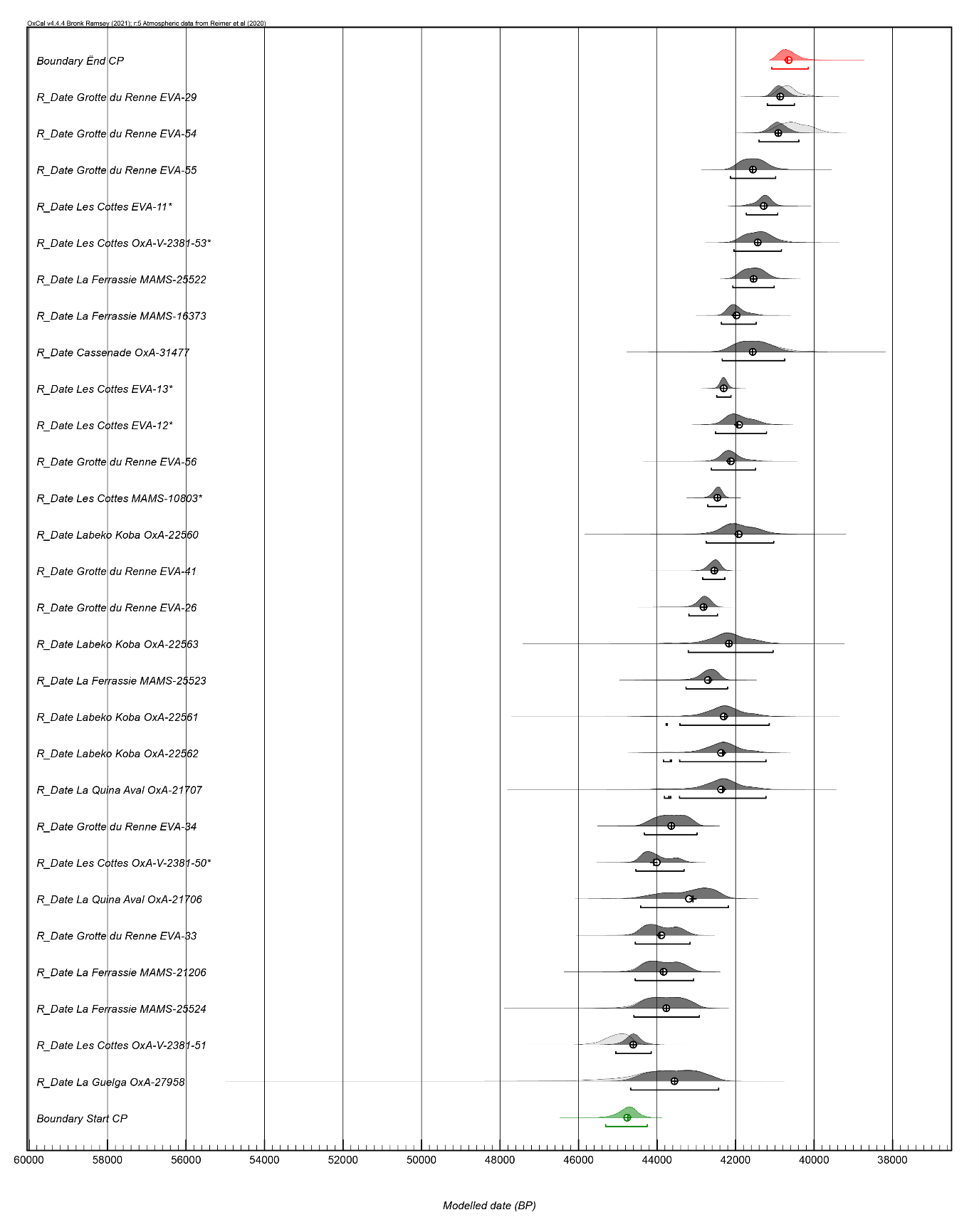

**Supplementary Figure S5.** Dataset of calibrated age ranges for directly-dated Neandertals (n=10) used for estimating the ‘extinction’ time of regional Neandertals with OLE modelling. The five lowermost individuals and the individual from Goyet (Q56-1) were dated with CSRA dating of hydroxyproline (Deviese et al., 2021). Produced in the OxCal software (v4.4) (Bronk Ramsey, 2006).

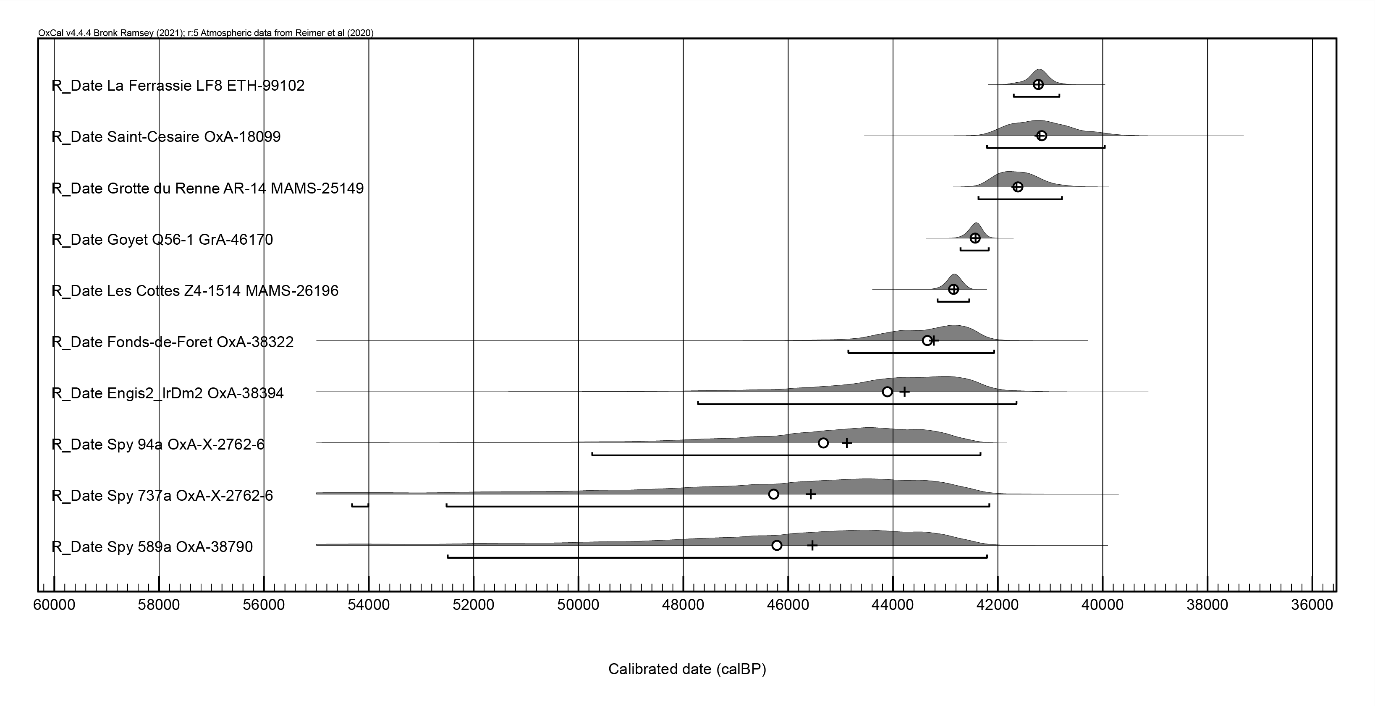

**Supplementary Figure S6.** Dataset of calibrated age ranges from Châtelperronian assemblages (n=8) used for estimating the ‘extinction’ time of the Châtelperronian with OLE modelling. All dates were prepared using ultrafiltration and dated using the AMS radiocarbon technique. Produced in the OxCal software (v4.4) (Bronk Ramsey, 2006).

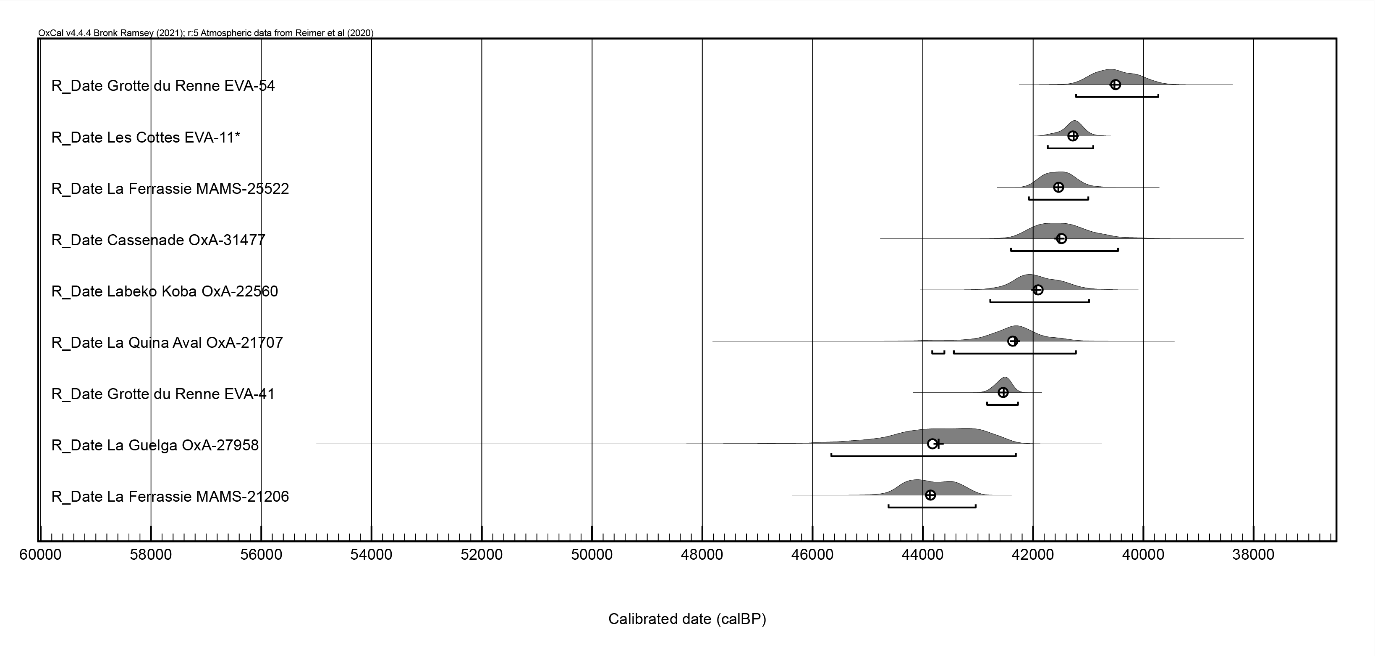

**Supplementary Figure S7.** Dataset of calibrated age ranges from Protoaurignacian assemblages (n=9) used for estimating the ‘emergence’ time of the Protoaurignacian with OLE modelling. All dates were prepared using ultrafiltration and dated using the AMS radiocarbon technique. Produced in the OxCal software (v4.4) (Bronk Ramsey, 2006). The earliest date from Isturitz (OxA-X-2694) was not included as discussed in the main text (see Barshay-Smidzt et al., 2018).

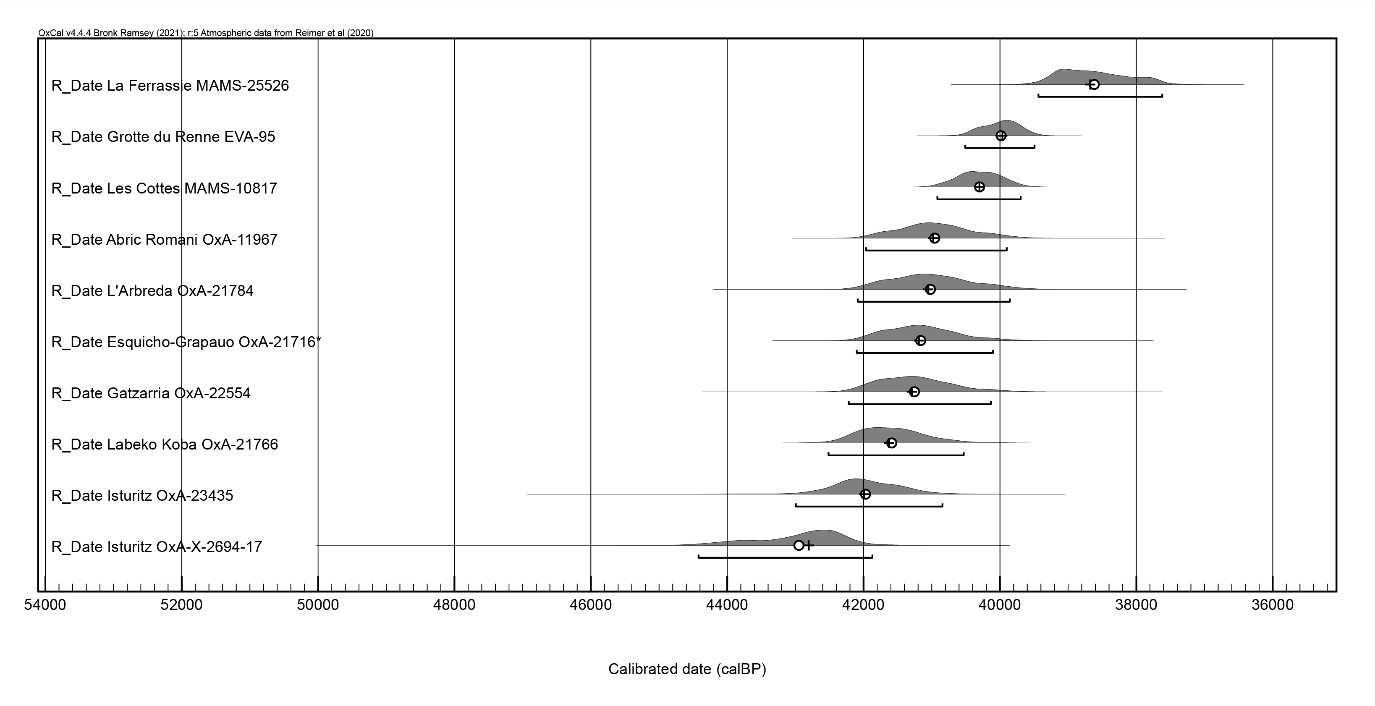

**Supplementary Figure S8.** OxCal script used to produce the kernel density estimation (KDE) plots and Bayesian start/end dates (**Figure 2**, main text).

**Plot()**

**{**

**Sequence()**

**{**

**Boundary("Start CP")**

**{**

**color="green";**

**};**

**Phase()**

**{**

**KDE_Plot("Chatelperronian")**

**{**

**};**

**Sum("Sum CP")**

**{**

**};**

**R_Date("La Guelga OxA-27958",40300,1200);**

**R_Date("Les Cottés OxA-V-2381-51",42360,370);**

**R_Date("La Ferrassie MAMS-25524",40770,650);**

**R_Date("La Ferrassie MAMS-21206",40890,500);**

**R_Date("Grotte du Renne EVA-33",40970,424);**

**R_Date("La Quina-Aval OxA-21706",39400,1000);**

**R_Date("Les Cottés OxA-V-2381-50*",41070,300);**

**R_Date("Grotte du Renne EVA-34",40520,389);**

**R_Date("La Quina-Aval OxA-21707",38100,900);**

**R_Date("Labeko Koba OxA-22562",38100,900);**

**R_Date("Labeko Koba OxA-22561",38000,900);**

**R_Date("La Ferrassie MAMS-25523",39000,510);**

**R_Date("Labeko Koba OxA-22563",37800,900);**

**R_Date("Grotte du Renne EVA-26",39390,334);**

**R_Date("Grotte du Renne EVA-41",38730,333);**

**R_Date("Labeko Koba OxA-22560",37400,800);**

**R_Date("Les Cottés MAMS-10803*",38540,270);**

**R_Date("Grotte du Renne EVA-56",37710,533);**

**R_Date("Les Cottés EVA-12*",37360,610);**

**R_Date("Les Cottés EVA-13*",38100,210);**

**R_Date("Cassenade OxA-31477",36660,750);**

**R_Date("La Ferrassie MAMS-16373",37380,390);**

**R_Date("La Ferrassie MAMS-25522",36590,390);**

**R_Date("Les Cottés OxA-V-2381-53*",36410,450);**

**R_Date("Les Cottés EVA-11*",36230,210);**

**R_Date("Grotte du Renne EVA-55",36630,452);**

**R_Date("Grotte du Renne EVA-54",35380,390);**

**R_Date("Grotte du Renne EVA-29",35500,216);**

**};**

**Boundary("Ënd CP")**

**{**

**color="red";**

**};**

**};**

**};**

**Plot()**

**{**

**Sequence()**

**{**

**Boundary("Start PA")**

**{**

**color="green";**

**};**

**Phase()**

**{**

**KDE_Plot("Protoaurignacian")**

**{**

**};**

**Sum("Sum PA")**

**{**

**};**

**R_Date("Isturitz OxA-X-2694-17",38900,1000);**

**R_Date("Isturitz OxA-23435",37500,900);**

**R_Date("Isturitz OxA-23436",37400,900);**

**R_Date("Isturitz OxA-23434",37000,800);**

**R_Date("Labeko Koba OxA-21766",36850,800);**

**R_Date("Labeko Koba OxA-X-2314-43",36500,750);**

**R_Date("Isturitz OxA-23432-33",37000,566);**

**R_Date("Gatzarria OxA-22554",36300,700);**

**R_Date("Esquicho-Grapaou OxA-21716*",36150,626);**

**R_Date("L'Arbreda OxA-21784",36000,700);**

**R_Date("L'Arbreda OxA-21665",35850,700);**

**R_Date("L'Arbreda OxA-21664",35900,650);**

**R_Date("Abric Romaní OxA-11967",35900,600);**

**R_Date("Esquicho-Grapaou OxA-21717",35550,750);**

**R_Date("Labeko Koba OxA-21793",35400,650);**

**R_Date("Trou de la Mère Clochette OxA-19622",35460,250);**

**R_Date("Les Cottés MAMS-10817",35150,280);**

**R_Date("Les Cottés OxA-V-2382-47",34870,340);**

**R_Date("Les Cottés MAMS-10816",34620,390);**

**R_Date("Grotte du Renne EVA-95",34810,210);**

**R_Date("Gatzarria OxA-22553",33800,550);**

**R_Date("L'Arbreda OxA-21674",33800,550);**

**R_Date("Les Cottés S-EVA-13669",34430,180);**

**R_Date("Les Cottés MAMS-10827",34080,250);**

**R_Date("Grotte du Renne EVA-81",33850,311);**

**R_Date("Trou de la Mère Clochette OxA-19622",33750,350);**

**R_Date("La Ferrassie MAMS-25526",33730,290);**

**R_Date("Les Cottés MAMS-10826",33710,230);**

**};**

**Boundary("Ënd PA")**

**{**

**color="red";**

**};**

**};**

**};**

**Plot()**

**{**

**Sequence()**

**{**

**Boundary("Start NDT")**

**{**

**color="green";**

**};**

**Phase()**

**{**

**KDE_Plot("Neanderthals")**

**{**

**};**

**Sum("Sum NDT")**

**{**

**};**

**R_Date("Saint-Césaire OxA-18099",36200,750);**

**R_Date("La Ferrassie LF8 ETH-99102",36170,220);**

**R_Date("Grotte du Renne AR-14 MAMS-25149",36840,660);**

**R_Date("Goyet Q56-1 GrA-46170",38440,320);**

**R_Date("Fonds-de-Forêt P44560 OxA-38322",39500,1100);**

**R_Date("Les Cottés Z4-1514",39485,271);**

**R_Date("Engis 2_LRdm2 OxA-38394",39900,1700);**

**R_Date("Spy 94a M3 OxA-X-2762-21",41500,1800);**

**R_Date("Spy 589a URd1 OxA-38790",41700,2300);**

**R_Date("Spy 737a OxA-X-2762-6",41600,2400);**

**};**

**Boundary("Ënd NDT")**

**{**

**color="red";**

**};**

**};**

**};**

**Supplementary Table S1.**

List of sites, countries of location, and number of dates (per category, site, and total) included in the study. All dates are produced after the year 2000 on osseous material and were dated using AMS ultrafiltration or HYP-CSRA.

| Site | Country | Number of Châtelperronian Dates | Number of Protoaurignacian Dates | Number of Neandertal (Direct) Dates | Dates total | References |
| --- | --- | --- | --- | --- | --- | --- |
| La Güelga | Spain | 1 |  |  | **1** | Higham et al., 2014 |
| Abric Romaní | Spain |  | 1 |  | **1** | Camps and Higham, 2012 |
| L'Arbreda | Spain |  | 4 |  | **4** | Wood et al., 2014 |
| Labeko Koba | Spain | 4 | 3 |  | **7** | Higham et al., 2014; Wood et al., 2014 |
| Cassenade | France | 1 |  |  | **1** | Discamps et al., 2020 |
| Grotte du Renne | France | 8 | 2 | 1 | **12** | Hublin et al., 2012; Welker et al., 2016 |
| La Ferrassie | France | 5 | 1 | 1 | **8** | Talamo et al., 2020; Balzeau et al., 2020 |
| La Quina-Aval | France | 2 |  |  | **2** | Higham et al., 2014 |
| Les Cottés | France | 7 | 6 | 1 | **14** | Talamo et al., 2011; Jouen et al., 2019 |
| Isturitz | France |  | 5 |  | **5** | Barshay-Szmidt et al., 2018 |
| Esquicho-Grapaou | France |  | 2 |  | **2** | Barshay-Szmidt et al., 2020 |
| la Mere Clochette | France |  | 2 |  | **2** | Szmidt et al., 2010 |
| Gatzarria | France |  | 2 |  | **2** | Barshay-Szmidt et al., 2012 |
| Saint-Césaire | France |  |  | 1 | **1** | Hublin et al., 2012 |
| Fonds-de-Forêt | Belgium |  |  | 1 | **1** | Deviese et al., 2021 |
| Spy | Belgium |  |  | 3 | **3** | Deviese et al., 2021 |
| Engis | Belgium |  |  | 1 | **1** | Deviese et al., 2021 |
| Goyet | Belgium |  |  | 1 | **1** | Rougier et al., 2016 |
| *TOTAL* |  | ***28*** | ***28*** | ***10*** | ***66*** |  |

**Supplementary Table S2.**

Dataset of calibrated radiocarbon ages used to estimate the start date of the Protoaurignacian in France and northern Spain using OLE modelling (*indicates the oldest date used in this model).

| Site | Lab code | cal BP 95.4% (IntCal20) | Reference |
| --- | --- | --- | --- |
| La Ferrassie | MAMS-25526 | 39435-37620 | Talamo et al., 2020 |
| Grotte du Renne | EVA-95 | 40509-39493 | Hublin et al., 2012 |
| Les Cottés | MAMS-10817 | 40918-39696 | Talamo et al., 2011 |
| Abric Romaní | OxA-11967 | 41961-39900 | Camps and Higham, 2012 |
| L'Arbreda | OxA-21784 | 42087-39855 | Wood et al., 2014 |
| Esquicho-Grapaou | OxA-21716 and OxA-21732 | 42101-40101 | Barshay-Szmidt et al., 2020 |
| Gatzarria | OxA-22554 | 42218-40133 | Barshay-Szmidt et al., 2012 |
| Labeko Koba | OxA-21766 | 42514-40531 | Wood et al., 2014 |
| Isturitz | OxA-23435 | 42994-40843* | Barshay-Szmidt et al., 2018 |

**Supplementary Table S3.**

Dataset of calibrated radiocarbon ages used to estimate the end date of the Châtelperronian with OLE modelling (*indicates the youngest date used in this model).

| Site | Lab code | cal BP 95.4% (IntCal20) | Reference |
| --- | --- | --- | --- |
| La Güelga | OxA-27958 | 45660-42312 | Higham et al., 2014 |
| La Quina-Aval | OxA-21707 | 43830-41224 | Higham et al., 2014 |
| Grotte du Renne | EVA-41 | 42836-42274 | Hublin et al., 2012 |
| Labeko Koba | OxA-22560 | 42777-40986 | Wood et al., 2014 |
| Cassenade | OxA-31477 | 42401-40458 | Discamps et al., 2020 |
| La Ferrassie | MAMS-25522 | 42076-41000 | Talamo et al., 2020 |
| Les Cottés | EVA-11 and OxA-V-2381-53 | 41735-40910 | Talamo et al., 2011 |
| Grotte du Renne | EVA-54 | 41221-39729* | Hublin et al., 2012 |

**Supplementary Table S4.**

Dataset of calibrated radiocarbon ages used to estimate the ‘extinction’ date of regional Neandertals using OLE modelling (*indicates the youngest date used in this model).

| Site | Lab code | cal BP 95.4% (IntCal20) | Reference |
| --- | --- | --- | --- |
| Spy 506a | OxA-X-2762-6 | 52518-42162 | Deviese et al., 2021 |
| Spy 92b | OxA-38790 | 52496-42207 | Deviese et al., 2021 |
| Spy 94a | OxA-X-2762-21 | 49738-42332 | Deviese et al., 2021 |
| Engis | OxA-38394 | 47721-41645 | Deviese et al., 2021 |
| Fonds-de-Forêt | OxA-38322 | 44216-41905 | Deviese et al., 2021 |
| Les Cottés Z4-1514 | MAMS-26196 | 43150-42545 | Jouen et al., 2019 |
| Goyet Q56-1 | GrA-46170 | 42713-42176 | Rougier et al., 2016 |
| Grotte du Renne AR-14 | MAMS-25149 | 42370-40778 | Welker et al., 2016 |
| La Ferrassie LF8 | ETH-99102 | 41696-40827 | Balzeau et al., 2020 |
| Saint-Césaire | OxA-18099 | 42206-39960* | Hublin et al., 2012 |
